# A scaling law for epithelial tissue rheology

**DOI:** 10.1101/2023.09.12.557407

**Authors:** M. I. Cheikh, N. Rodriguez, K. Doubrovinski

## Abstract

Epithelial morphogenesis is a process through which simple cellular sheets are shaped into complex tissues and organs in a developing animal. From a physics perspective, understanding any shape change requires knowing the active forces driving its dynamics, as well as the material properties, i.e. the rheology, of the material that is undergoing the deformation. Despite a long-standing effort, rheological properties of embryonic tissues have remained elusive. Here, we develop a minimal theory providing a comprehensive explanation of rheological measurements characterizing the mechanics of epithelia in the early fly embryo. Our theory explains a key experimental observation: when subjected to concentrated pulling force, the embryonic epithelium of the fruit fly *Drosophila melanogaster* deforms following a power law with an exponent of 1/2. All dimensional parameters of our theory are constrained by direct measurements and have allowed us to estimate the spring constant of an individual cellular edge. We show that stress relaxation (attributable to actin turnover), stretching elasticity of individual cellular edges, and the floppy topology of the cellular network are the sole physical properties governing tissue rheology on the developmentally relevant time scale of 1-10 minutes.

---

In the recent years, it has been increasingly appreciated that morphogenesis is fundamentally a mechanical process, with numerous examples of computational models proposed to explain the physical mechanisms underlying tissue mechanics [1–6]. While these models have in many cases been successful at describing known morphogenetic phenomena, it is often unclear whether they have accurately depicted the underlying physics, and therefore it is also unclear whether they can be used to make meaningful predictions. Arguably, the reasons for this are mainly two-fold. In most cases, the relevant parameters have not been experimentally measured, so there is the possibility of over-fitting; see [7] for an excellent example demonstrating this point. More broadly, it is unclear which particular physical effects (e.g. elasticity, viscosity, active stresses, cellular geometry, etc.) play key roles in tissue morphogenesis.

The embryo of the fruit fly *Drosophila melanogaster* is one of the most prominent model systems for studying epithelial morphogenesis. Physical models have been proposed for developmental processes including ventral furrow formation [8], germ band extension [9], and dorsal closure [10, 11]. Only recently has work been done to experimentally measure material properties of these tissues, and only at early developmental stages [12–17]. The early *Drosophila* embryo is an ellipsoid tissue in which a single monolayer of epithelial cells encases a viscous [15, 16] yolk-filled interior (Fig. 1). Recent work has demonstrated that the epithelial portion of the embryo is largely elastic [16], that this elasticity is due mostly to cortical (cell-surface-associated) actin [16], and that the basal portion of the cells (which have extremely high concentrations of actin) overwhelmingly dominate elasticity [17]. Further experiments have demonstrated that external friction between the tissue and the surrounding eggshell is negligible ([17], but see [18]), that active myosin-generated stresses have negligible effect on passive material properties [19], and that minute-time-scale ventral furrow dynamics occurs in the adiabatic regime [17]. Based on this information, we have been able model the behavior of the embryo both in response to externally applied forces [17] and in response to the endogenous myosin-generated forces that drive ventral furrow formation [20]. However, a key observation we have not been able to explain is that deformation due to a constant externally applied force obeys the power law *X*(*t*) *∼ t*^1*/*2^, where *X* is the displacement of the point being pulled and *t* is time [16, 19]. In general, power laws reflect general features of the dynamics, and are insensitive to model details such as the specific values of the dimensional model parameters. Specific scaling exponents characterizing mechanical properties of individual cells (see e.g. [21, 22]) and the spreading dynamics of cells on substrates [23, 24] have been reported in the past. Here, we develop a consensus theory explaining the power law scaling for epithelium described above [16, 19]. As a confirmation, our description provides an estimate of the elasticity of individual cellular edges which agrees closely with our previous independent estimate.

**FIG. 1.**
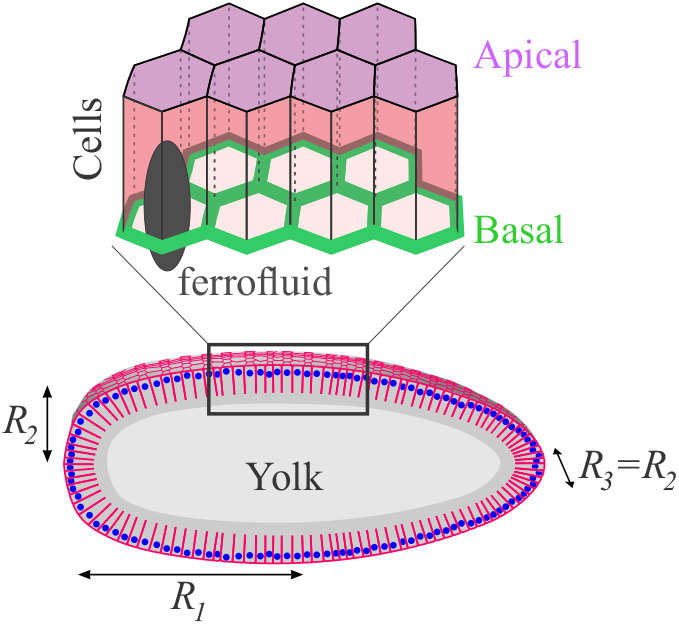
Schematic geometry of the early embryo of the fruit fly *Drosophila melanogaster*.

The paper is organized as follows. First, we briefly review the relevant experimental setup. Next, we provide additional data strengthening some of our previous experimental findings. Thereafter, we describe a model of epithelial tissue rheology that provides a minimal quantitative description of our data. Next, to give physical intuition behind the conclusions, we formulate a simplified 1d model, capturing the essential features of the full description. Finally, we use our minimal model to deduce the elastic spring constant of cellular edges in the early fly embryo and compare the obtained value to our previous independent measurement. The implications of our results are further discussed in the concluding section.

Fig. 2a illustrates the experimental setup (see also Fig. 1 for an explanation of embryo geometry). First, an embryo is injected with a droplet of ferrofluid. Next, one exploits an unusual feature of *Drosophila* development: at the blastula stage, cells are open on one side. In effect, each cell has a small hole on the basal side, through which materials including cytoplasm and organelles can pass freely without mechanical hindrance. This feature of fly development allows one to force the ferrofluid droplet into the interior of one cell by carefully controlling its position using a permanent magnet mounted next to the embryo. Once the ferrofluid droplet has been positioned within a cell, the magnet is moved away from the droplet such that the droplet starts moving towards the magnet along the surface of the embryo, stretching the cellular layer as it moves. Since droplet displacement during the course of an experiment is small compared to the distance between the droplet and the magnet, the force on the droplet may be assumed to be approximately constant (as was experimentally verified in [16]). In these experiments, it was shown that the displacement of the droplet subjected to constant force increases according to a power law with an exponent of 1*/*2. To test these results further, here we have performed additional experiments where the time during which the force is applied to the tissue is increased between three-to six-fold compared to previously published data (Fig. S1). Additionally, we performed measurements using an electromagnet setup to further improve signal to noise (Fig. 2b.) Time-displacement curves can readily be collapsed on a master curve following a 1*/*2 power law (Figs. 2b, S1) in agreement with the earlier work [16, 19].

**FIG. 2.**
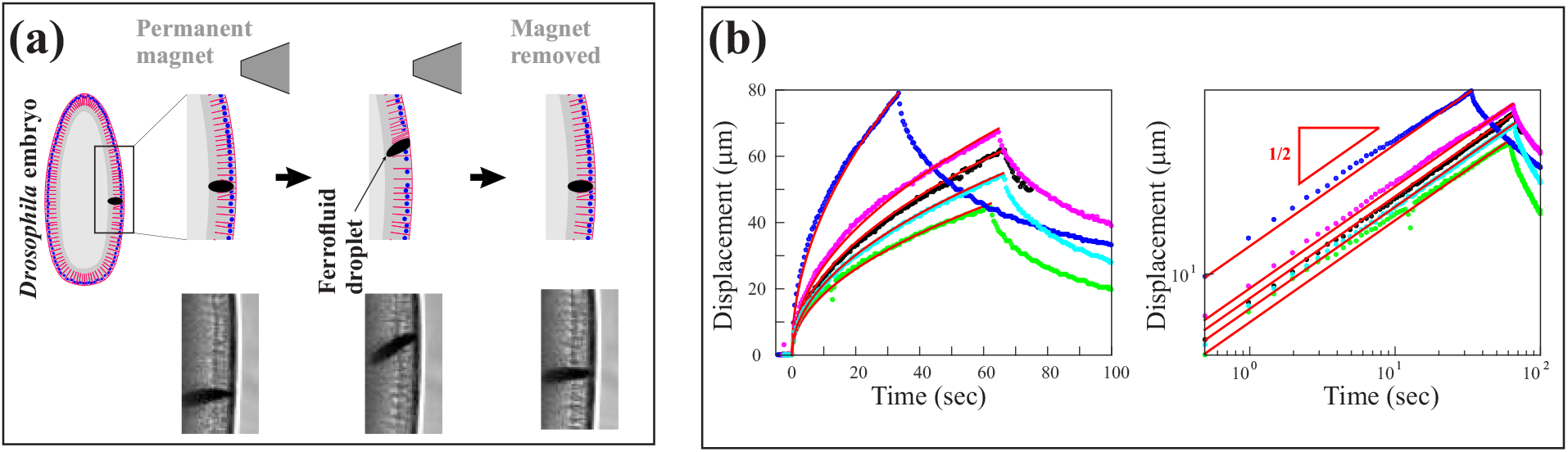
Power law response in tissue pulling experiments. (a) Schematic explanation of experimental setup. Top panel illustrates an experiment where a ferrofluid droplet is first positioned in the interior of one cell, and then used to stretch the epithelium by an externally applied magnetic field. Bottom panel illustrates snapshots from an *in vivo* measurement. (b) Left panel: when a ferrofluid droplet inserted into the cellular layer is pulled with constant force, displacement scales like time to the power 1*/*2 (0.48 *±* 0.02, mean *±* standard error, n=5). The force was applied for approx. 60 seconds (see Supplement for measurements over a longer time range). Right panel shows the same set of traces on a log-log plot.

To understand the physical nature of the observed scaling, we developed a minimal model of epithelial tissue. We describe cells as hexagons, whose edges are linearly elastic springs with spring constant *k* and a dynamic rest length (see below). It was previously shown that elasticity of the embryo is completely dominated by its actin cytoskeleton, which turns over on a time scale of tens of seconds [25]. Actin is exclusively associated with the cellular boundaries [16, 25], and actin specifically on the basal sides of cells makes up the overwhelming contribution to the total effective elasticity of the tissue [16, 17]. Basal sides of cells make up a continuous hexagonal network encasing the yolk (Fig. 1). Since tissue elasticity is dominated by this structure, our minimal model will only represent basal sides of cells. The apical and lateral sides are not described explicitly and are only assumed to prevent out-of-plane deformation of the basal side when it is subjected to the pulling force. We therefore describe the tissue as a hexagonal network of linearly elastic elements, whose vertices are confined to the surface of an ellipsoid. The three semi-axes of the ellipsoid will be denoted by *R*_1,2,3_. We set their ratios to the experimentally relevant values of *R*_1_*/R*_2_ *≈* 3, *R*_2_*/R*_3_ = 1, and do not vary those ratios in what follows. Δ will denote the initial (mean) length of an individual network edge, representing basal edge of a cell. Rest lengths of cell edges are dynamic, relaxing to their respective instantaneous lengths on a timescale *τ* :

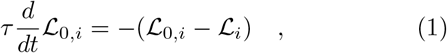

where ℒ_*i*_(*t*) is the length of the *i* :th cell edge, *τ* is the time-scale of elastic stress relaxation, and ℒ_0,*i*_(*t*) is its instantaneous rest length (in particular, ℒ_*i*_(*t* = 0) *≈* Δ). The area of individual cells comprising the mesh was not constrained, since cell volume rather than area is conserved in this tissue [16, 17]. To simulate external pulling force from a ferrofluid droplet, a constant force of magnitude *F* is applied to one vertex of the cellular network. Dynamics of each cell vertex is given by the force balance

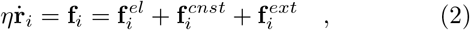

where **r**_*i*_ is the position of the *i*:th vertex of the mesh, and *η* is effective friction coefficient, assumed negligible in what follows. **f**_*i*_ is total force on the *i* :th vertex comprising the following contributions: (1) elastic force from adjacent cell edges 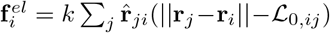, where the sum is over all edges adjacent to the *i*:th vertex, ℒ_0,*ij*_ is the rest length of the edge between vertices *i* and *j*, and 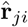 is the unit vector along that edge, (2) force of constraint 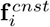 confining the vertex to the surface of the ellipsoid (normal to ellipsoid surface and directed towards the surface; its magnitude is chosen to dominate any other force contribution), and (3) force 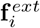 of magnitude *F* due to external pulling. The last contribution is only applied to a single vertex assumed to be in contact with the ferrofluid droplet. Finally, the node being pulled interacts with edges through “excluded volume” interactions such that it drags along with it any edges that it encounters in its path. In all simulations, the hexagonal mesh representing basal cell edges is approximately uniform, such that the lengths of individual edges do not deviate significantly from the mean; we checked that all our conclusions are robust with respect to the degree of disorder within the mesh. Rescaling time, length and force with *τ*, Δ, and *F* respectively, we are left with the following dimensionless parameters: elasticity of a cellular edge *k*Δ*/F*, effective friction with the viscous environment *η*Δ*/*(*Fτ*), and the three semi-axes of the embryo *R*_1,2,3_*/*Δ.

In our simulations, when subjected to constant pulling force, the displacement of the vertex that is being pulled follows a 1*/*2 power law, in agreement with previous experimental findings [16, 19], see Fig. 3b (see also Fig. 4 for a log-log plot demonstrating the scaling).

**FIG. 3.**
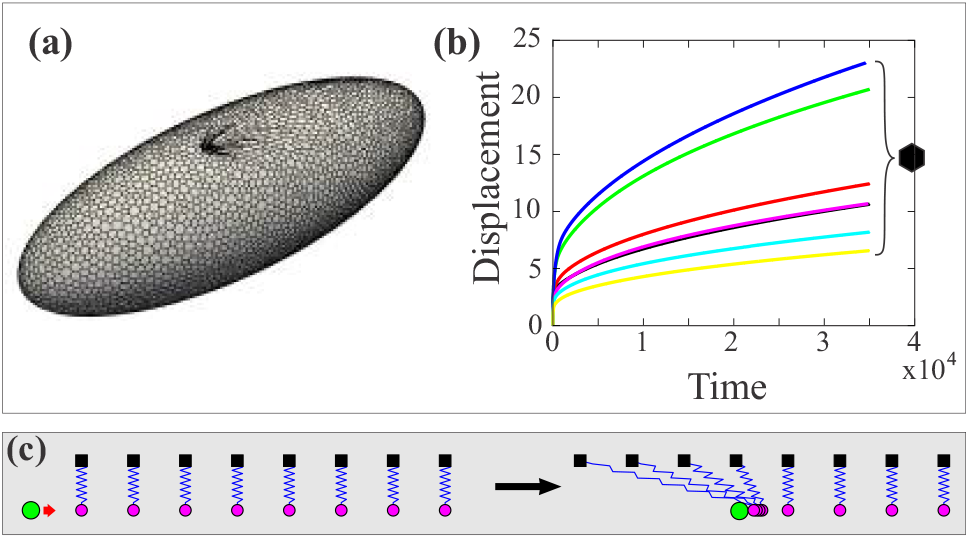
Simulations of the minimal model reproducing the observed scaling. a). Simulated system is a network of hexagons, confined to a surface of an ellipsoid that is being stretched by concentrated force (see also Fig. S3 for a representative plot of the displacement field). b). Simulations of the system in a). Curves show displacement of the point being pulled as a function of time. See Supplementary Material for simulation details. c). Schematic of the 1D simplified model. A point (green circle) moves through a lattice of springs, engaging with one end of each spring as it moves past it. Note that springs are depicted as being vertical for illustration purpose: in our actual description forces only have a horizontal component.

**FIG. 4.**
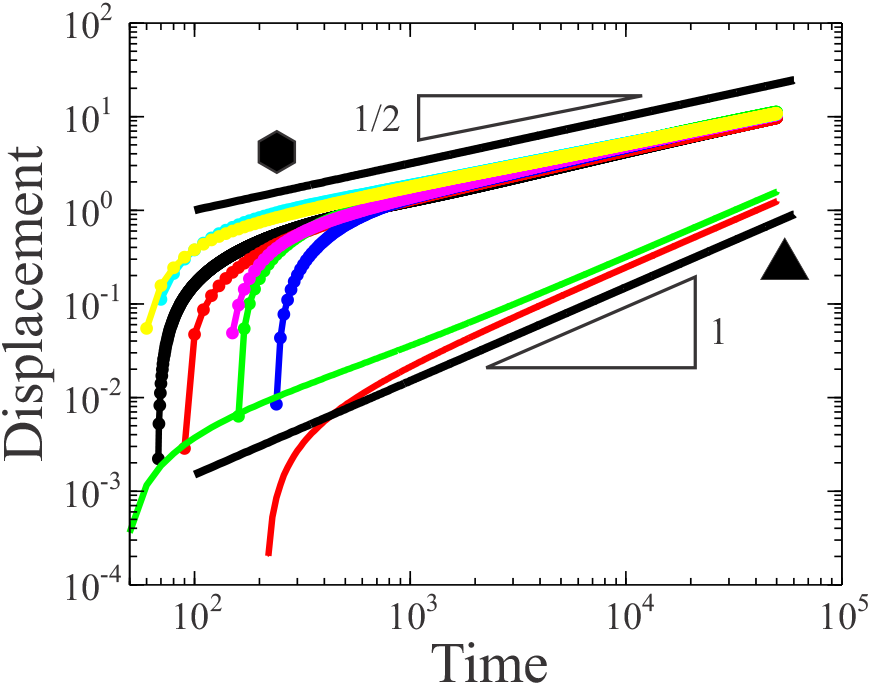
Re-scaling confirming that the simplified model correctly explains the scaling behavior in the full equations. Two curves in the bottom of the figure show simulations of triangular meshes. See Supplementary Material for simulation details.

To understand the origin of the power law scaling, we now turn to simulations representing limits of the original system. First, instead of considering a network of hexagons, consider a network of triangles. In this case, the asymptotic displacement follows a power law with an exponent of 1, and not 1*/*2 (Fig. 4). A network of triangles is a good approximation of a linearly elastic sheet [26, 27], which effectively behaves as a single elastic spring. Response of a single spring “forgetting” its rest-length on a finite time-scale is asymptotically viscous. Thus, asymptotically, constant force results in constant speed of displacement, in other words, a power law with an exponent of 1.

Why do we recover an exponent of 1*/*2 in the case of hexagons? A hexagonal mesh is an example of a floppy network, meaning, in particular, that some deformation can be accommodated without energy cost. Specifically, the transition between floppy and stiff rheology is governed by coordination number *z*, the number of bonds per node [28]. In a triangular network, every node is adjacent to six edges, i.e. *z* = 6, whereas *z* = 3 for a hexagonal mesh. Simple counting shows that for the number of degrees of freedom in the system to exceed the number of constraints one must have *z >* 2*d*, where *d* is the spatial dimension, i.e. *z* = 4 in our case. As a result, networks of hexagons are floppy, whereas networks of triangles are stiff. From this, informally, it may appear plausible that hexagons and triangles show qualitatively different behaviors, but it is not clear why we specifically recover the exponent of 1*/*2. Floppy network have recently been the focus of much research, see in particular [28, 29], as well as more recent literature. However, to the best of our knowledge, this research has mainly focused on the rigidity-percolation transition and the associated critical behaviors, whereas large deformations and finite size effects, although very relevant to *in vivo* data, have received less attention.

Why do we then recover a power law of 1*/*2 in the case of hexagons? A simple physical picture (tentatively) capturing the observed behavior in the floppy case is as follows. First, consider a hexagonal network of springs whose rest lengths are not dynamic. As the ferrofluid droplet moves through such a network, it encounters more and more elastic elements that it drags along. This is in stark contrast with the linearly elastic network (of triangles) where the entire network engages in the deformation from the beginning of pulling. Indeed, linearity implies that displacement at any given point is proportional to the displacement of the point being pulled. Roughly, in the linear case of triangles, as the droplet moves through the network, all springs stretch more and more. In the floppy case, instead, more and more springs stretch.

To test our simplified physical picture, we analyzed a simplified 1D model meant to capture the conjectured behavior. Consider a 1D lattice of springs, each of initial length zero (Fig. 3c). Consider a point (or droplet), moving through the spring lattice. As it encounters springs in its path, it engages with one end of the spring, the other end remaining in place, such that the spring stretches out as the point moves past it. Additionally, rest lengths of those springs are dynamic, relaxing on a finite time-scale *τ*. As shown in the Supplementary Material, the asymptotic solution to this 1D model is given by:

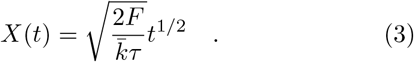

That is, our simplified 1D model predicts the 1*/*2 power law scaling. To check whether the 1D model captures the relevant features of the original 3D network of hexagons, we next mapped the parameters of the full system to those of the simplified 1D model. Using geometric arguments (see Supplemental Material), we set *τ ∼ τ*_1*d*_, *F ∼ F*_1*d*_, and 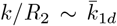, where the 1*d*-subscript denotes the parameter of the 1D model.

Combining the scaling assumptions with Eq. 3 gives:

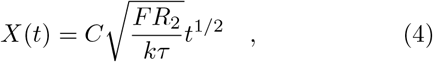

where *C* is a dimensionless shape-factor independent of model parameters.

If the analogy between the simplified 1D model and the full description indeed holds, rescaling the curves from Fig. 3b by 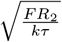 must collapse all those curves on one master-curve. Fig. 4 shows that this is indeed the case. This is strong evidence that Eq. 4 accurately represents the key aspects of the dynamics. Note that the *t*^1*/*2^ power law breaks down at small time due to finite-size effects. Clearly, this must be the case, since otherwise the velocity would be infinite at *t* = 0. Experimental data corresponding to small times is not very reproducible and we do not expect these to be accurately explained by a simplified model not taking finite size effects into account (e.g. the ferrofluid droplet has a finite size; the droplet changes its shape when the magnetic field is applied, influencing the precise details of the measured curves at very short times, etc). In Fig. S4 we simulated the case where the tissue had a large transverse cut behind the point being pulled. This did not qualitative alter the dynamics, suggesting that only accounting for elastic elements in front of the point being pulled is an adequate approximation.

To further test our theory, we varied the magnitude of the externally applied force. This may be done by using the electromagnet setup and varying the current in the coil (see Supplemental text and Fig. S5 for the details of the setup). We found that a change in the magnitude of force by a factor of 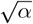, resulted in the displacement of the droplet as a function of time to increase by a factor of *α*, in complete agreement with our scaling theory in Eqs. 3,4, lending further support to our main conclusions (Fig. S5). Additionally, we attempted to block actin turnover pharmacological to see if the 1*/*2 scaling would turn into exponential relaxation. We were only partially successful, managing to block turnover within large tissue patches, but not across the entire embryo (Fig. S6). The corresponding pulling experiments decreased the amplitude of the displacement curves, but did not affect the power law exponent, in agreement with the corresponding simulations (Fig. S6).

A fully quantitative comparison would require parameter-free comparison between the pre-factor measured in experiments, and that in Eq. 4. From simulations we obtain *C ≈* 36. The external force *F* in our measurements from [16] was calibrated with *F ≈* 20 nN. In recent measurements from the lab of Dr. Anna Sokac it was shown that 80 % of F-actin comprising basal cell edges turns over on a time-scale of approx. 20 sec [25], implying *τ ≈* 20 sec. The radius of the embryo *R*_2_ is approx. 100 *μ*m. The only unknown parameter left is *k*, the spring constant of a cellular edge. Experimental curves in Fig. S1 match parabolas exactly, with no significant systematic deviation, as is immediate from Figs. 2b, S1. Fitting these parabolas to expression 4, using the measured values for *F, τ*, and *R*_2_, we estimate *k ≈* 23 *±* 8.9 *nN/μm* (mean standard *±* error, n=5), which agrees well with our independent estimate of *k* = 6.8·5 = 34 *nN/μm* from [17] (see in particular the last row in the table of Fig. 7c” therein). The factor of 5 comes from the fact that the value obtained in [17] is relevant for the stable fraction of actin that does not turn over and comprises 20 % of the total pool.

Arguably, the major limitation in formulating quantitative models of morphogenesis is limited knowledge of precisely which physical effects dominate the rheology of animal epithelia. Here, for the first time, we have been able to develop a quantitative and (practically) parameter-free physical description of a biological tissue, relating properties of cells to the rheology of the tissue. We have shown that elasticity of cellular edges, stress relaxation (attributable to actin turnover), and the floppy topology of the network are key physical aspects governing tissue rheology on the timescale of minutes (directly relevant to morphogenesis in the early embryo). Importantly, these results argue that bending contributions are not significant in this epithelium, since bending contributions would render a floppy hexagonal network stiff [29]. By the same token, volume conservation within individual cells seems to also not play a significant role. This may not be surprising in our system, since cells are open on their basal side. However, we expect this to be true more generally, if extension along the apico-basal axis does not require significant energy cost (as was shown for some vertebrate cells in [4]). Note additionally that a continuum description, in particular linear elasticity, can not be used to describe our system since the theory of linear elasticity can not account for floppiness. Any description of the tissue based on such theory would err on the specific power laws and orders of magnitudes of the corresponding dimensional parameters. We believe that our work will provide a rigorous foundation for future investigations of epithelial tissue rheology in a key model system. Notably, it will be interesting to investigate how cellular rearrangements (proceeding through T1 transitions and cellular rosettes) affect tissue rheology on longer time scales during the later developmental stages [30].

## Supplementary Material

### I. ADDITIONAL MEASUREMENT DATA WITH EXTENDED TIME RANGE

Whereas Fig. 2 of the main text showed data where ferrofluid droplet was subjected to force for approximately 60 seconds, the time-period can be extended to approximately 200 seconds, indicating that the power law scaling persists over almost 3 decades in time, see Fig. S1.

**FIG. S1.**
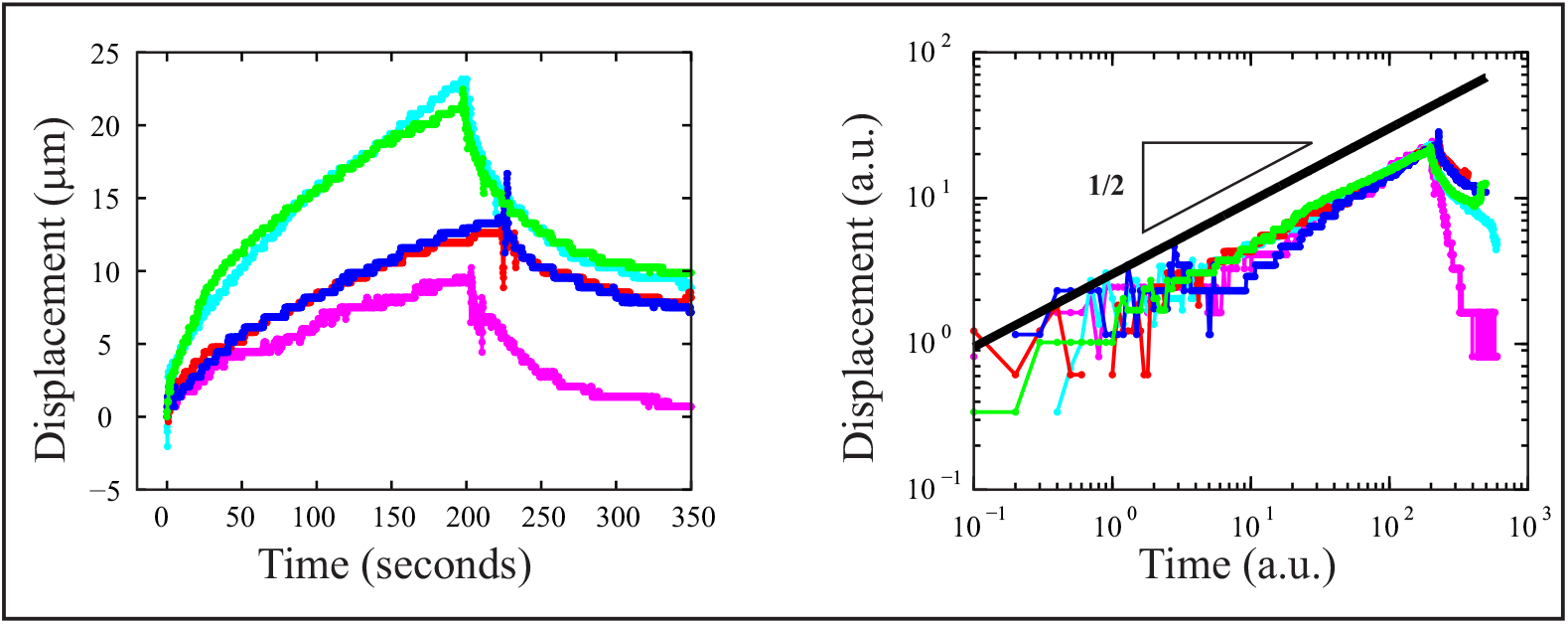
Power law relation tested over a longer time range. Experiments analogous to those in Fig. 2, except that force was applied for approx. 200 seconds, compared to approx. 60 sec in the previously reported experiments. These experiments were done using a permanent magnet, rather than the electromagnet used for Fig. 2. Right panel shows the same set of traces on a log-log plot.

### II. SIMULATIONS

All simulation code is publicly available at https://github.com/mcheikh1991/Results_Scaling_Law_For_Epithelial_Tissue_Rhelogy.git. For the hexagonal cases (Figs. 3 and 4), parameters are as follows: *k*Δ*/F* = 40, *η*Δ*/*(*Fτ*) = 0.2, ellipsoid half-axes *R*_1_*/*Δ and *R*_2_*/*Δ = *R*_3_*/*Δ are 32 and 11 (black), 42.7 and 15 (red) 94.2 and 33 (green), 126 and 44 (blue), 42 and 15 (magenta), 31 and 11 (cyan), 125 and 9 (yellow). For the triangular cases (Fig. 4), parameters are as follows: *k*Δ*/F* = 40, *η*Δ*/*(*Fτ*) = 200 (red), *η*Δ*/*(*Fτ*) = 20 (green), ellipsoid semi-axes *R*_1_*/*Δ and *R*_2_*/*Δ = *R*_3_*/*Δ are 63 and 22 respectively. In Fig. 4 a small offset was subtracted from the collapsed curves to minimize the influence of the finite size effects at short times on the scaling (the offset was the same for all seven curves; not subtracting off the offset worsens agreement slightly, putting the asymptotic slope at 0.4 instead of 0.5). Hexagonal mesh was generated using freeware Gmsh, all input files required to run the code are included in the git archive. Note also that the “hexagonal” mesh must contain some non-hexagonal cells to tile an ellipsoid.

### III. SIMPLIFIED 1D DESCRIPTION

As discussed in the main text, we analyzed a simplified 1D model meant to capture the conjectured behavior. Consider a 1D lattice of springs, each of initial length zero, see Fig. 3c. Consider a point (or droplet), moving through the spring lattice. As it encounters springs in its path, it engages with one end of the spring, the other end remaining in place, such that the spring stretches out as the point moves past it. Additionally, rest lengths of those springs are dynamic, relaxing on a finite time-scale *τ*. The dynamics of the system are:

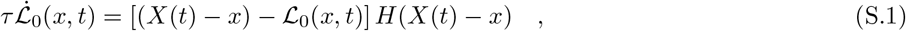

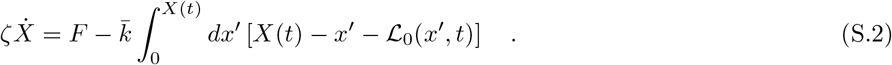

In Eq. S.1, ℒ_0_(*x*) is the rest length of spring at spatial position *x*, and *X*(*t*) is the position of the point pulled through the lattice. Initially, this point is at *x* = 0, i.e. *X*(0) = 0. Note that *X*(*t*) *− x* is the instantaneous length of the spring initially attached at position *x. H* is the Heaviside function, reflecting that only springs behind the ferrofluid droplet are stretched. Equation S.2 gives the dynamics of the point being pulled and follows from force balance comprising three contributions: (1) external pulling force *F*, (2) cumulative force from all springs encountered up until the time *t* (the integral term), and (3) effective viscous friction with the surrounding 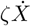. Note that Eqs. S.1, S.2 describe a 1D spring lattice in the continuum limit, such that 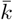 is defined as the density of elastic elements and has units of force per length squared.

To analyze Eqs. S.1, S.2, we integrate Eq. S.1 with respect to time, substitute the result into S.2, and interchange the order of integration in the second term on the r.h.s. to obtain

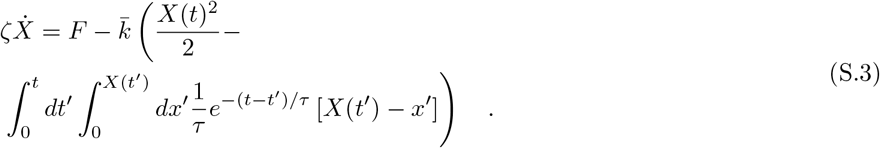

The integral term can be further simplified to 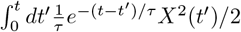. Substituting an ansatz *X*(*t*) *∼ t*^1*/*2^, dropping all terms that tend to zero asymptotically, and assuming *ζ* is negligible, we obtain the asymptotic solution to Eqs. S.1, S.2:

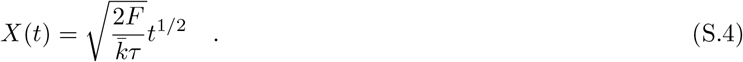

To summarize, our simplified 1D model predicts the 1*/*2 power law scaling, in agreement with simulations from Fig. 3.

### IV. MAPPING PARAMETERS FROM 1D TO 3D MODEL

To check whether this analysis applies to the original network of hexagons, we next formally map the parameters of the full system to parameters of the simplified 1D model (Eqs. S.1, S.2), making a specific set of assumptions about how the parameters of the full model scale with the parameters of the simplified description. In particular, we set *τ ∼ τ*_1*d*_, *F ∼ F*_1*d*_, and 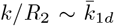, where the 1*d*-subscript denotes the parameter of the 1D model. To better understand the scaling assumption for the spring constant of an edge, consider the related case of a cylinder, whose radius is that same as *R*_2_, the radius of the model embryo, see Figs. 3a, S2. Suppose evenly spaced elastic encircle the cylinder, as in Fig. S2. These (fictitious) “rings” are meant to effectively capture the behavior of elastic elements comprising the floppy network in the original model and will correspond to individual springs in the simplified 1D description. These “rings” were inspired by a previous observation that the deformation field in an embryo subjected to localized pulling force resembled the stretching of (imaginary) ellipsoids encircling the short axis of the embryo (see Fig. Fig. 8d in [1] for further discussion). Additionally, similar displacement fields are seen in the full 3D model presented in this paper, Fig. S3. Note however, that these rings are just an effective construct (perhaps akin Edwards tubes in the theory of polymer melts) and we do not literally imply that such rings will indeed be apparent in the dynamics of the full system. Subdividing one such ring into segments of length Δ, each with spring constant *k*, the combined spring constant of one ring scales like *∼ k*Δ*/R*_2_. The density of these rings per unit length of the cylinder scales like *∼* 1*/*Δ. Multiplying these two expressions, we obtain *k/R*_2_ for the scaling of 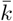 in Eq. S.4.

**FIG. S2.**
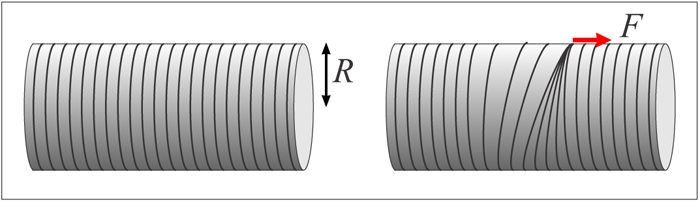
Schematic explaining the relation between the full model and the 1D approximation, see text.

**FIG. S3.**
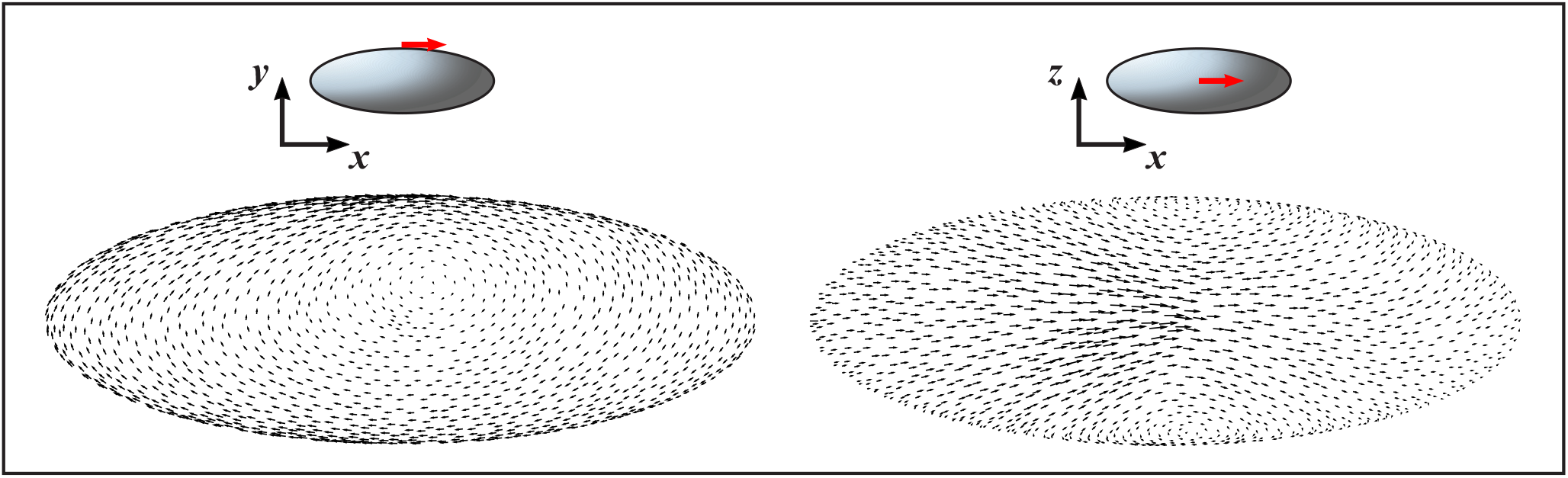
Displacement field during pulling. This represents the displacement field during pulling of one of our standard simulations (olive curve in S6c, and lime-green curve in Fig. S4. The arrows are scaled so as to not overlap. Left field shows view from the side (with pulling at the top, red arrow in schematic), right shows the same displacement field viewed from the top.

### V. FURTHER TEST OF SCALING THEORY BY SIMULATING A TRANSVERSE CUT

Our simplified 1D model (and our interpretation of the 1D and 3D models in Eqs. 3,4) suppose that elastic elements of the tissue in front of the point being pulled are responsible for the observed scaling. In other words, the elastic elements behind the point being pulled are assumed not to affect the scaling. To test this assumption, we simulated a situation where the tissue has a large cut right behind the point that is being pulled, see Fig. S4. It may be seen that the scaling is not significantly affected, although the displacement does increase somewhat, compared to the case without the cut. This result lends further support to our simplified theory described in the beginning of the Supplement.

### VI. SCALING WITH THE APPLIED EXTERNAL FORCE

An additional prediction of the theory is the square root scaling of the displacement curve with the applied force. Specifically, Eq. 4 implies that if the force is increased by factor *α*, the displacement of the droplet as a function of time is re-scaled by a factor 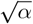. To test this prediction, it is necessary to repeat measurements with variable force and ensure that all other parameters are kept the same between measurements. To this end, we modified the setup described in our previous work [2]: instead of using permanent magnet as was done in our previous work, we used an electromagnet, as illustrated in Fig. S5. The use of electromagnet offers several key advantages. First and foremost, the force may be dialed precisely by dialing a specific value of the current. Second, the force is much stronger than that from a permanent magnet. This allows positioning the magnet much further away from the sample such that the field to which the ferrofluid droplet is subjected is almost completely constant during its motion. Finally, the onset of the force is controlled by a switch, such that the time-point of force application is sharply defined. Fig. S5 shows measurement results. In these experiments, the droplet was subjected to three consecutive force steps of increasing magnitude by dialing the coil current to 1, 2, and 3 A, and allowing the droplet to relax back to its initial position between the force steps. Re-scaling the measured curves by the inverse square root of the force (force being proportional to the current), it is seen that the data collapses onto a single curve within an excellent precision, lending further support to our scaling theory.

**FIG. S4.**
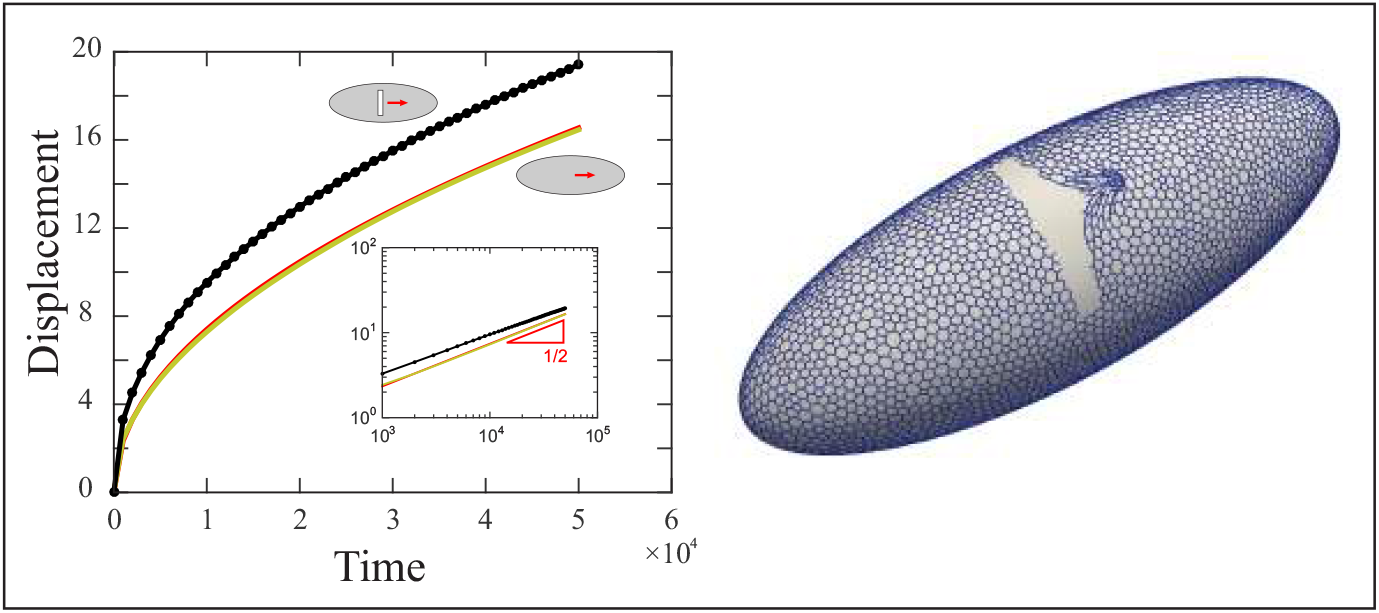
The scaling is not significantly affected if the tissue behind the point being pulled is separated from the region of force application by a large transverse cut. Left: lime-green curve is the original simulation of the case without the cut. Red curve (practically indistinguishable from the lime-green curve) is a fit to *t*^1*/*2^ power law. Black curve is simulation with the cut. Inset shows the same three curves on a log-log plot. Right panel is a snapshot of the 3D simulation.

### VII. EFFECTS OF BLOCKING ACTIN TURNOVER (WITHIN A CONFINED REGION)

To further test the validity of the proposed theory, we attempted to block actin turnover pharmacologically, to examine the effects of actin turnover on our measurements. To this end, we employed a protocol previously based on the work of Dr. Orion Weiner, utilizing a mixture of Rho kinase inhibitor Y-27632, myosin inhibitor blebbistatin, and an inhibitor of actin depolymerization jasplakinolide [3, 4]. (Additional experiments showed that Y-27632 and blebbistatin, included in the original protocol, were dispensable in this system.) Unfortunately, it proved impossible to block actin turnover in all of the tissue within an embryo. Specifically, when injected into the yolk sack, the drug cocktail becomes diluted and actin dynamics are not blocked efficiently. This may be overcome to an extent by injecting the cocktail into the narrow perivitelline space which separates embryonic epithelium from the rigid outer shell encasing the embryo (the vitelline membrane). We found that in this case, actin turnover was efficiently blocked within a portion of the embryo that could span as much as approximately 70 percent of embryo length and 50 percent of the width. In order to visualize actin dynamics, the corresponding experiments were done in a genetic background overexpressing an actin marker F-tractin-tdTomato (generated by crossing flies carrying the UASp-driven construct (Bloomington *Drosophila* Stock Center RRID:BDSC 58988) to maternal tubulin driver (RRID:BDSC 80361), see [5]). As may be seen from Fig. S6d, this resulted in appearance of intense red labeling in the vicinity of the injection site, indicating that actin turnover was efficiently blocked in that region. Additionally, this treatment completely blocked the progression of cellularization (gradual outgrowth of the membranes) within the injected region, as may be seen from the corresponding bright field image (Fig. S6d”). To visualize membranes, injection mixture also included CellMask dye (Invitrogen Cat. nr. C10046); membrane (rather than cytoplasmic) localization of dye indicates intact membranes (Fig. S6d’). In order to assess the effects of the treatment on the mechanical properties of the tissue, we followed the injection of the drug with injection of a ferrofluid droplet, and then positioned the droplet within the affected region. Next, we repeated pulling assay as in Fig. S1. Plotting displacement of the droplet as a function of time, it is seen that it follows the same 1*/*2 power law scaling as in the case of an untreated embryo (Fig. S6a,b). However, typical maximal displacement of the droplet appeared markedly decreased (note that these data should be compared to figures 1 and S8 in reference [2], whereas traces in Fig. S1 of the current paper show smaller maximal displacement since the magnitude of the applied force had to be set smaller to achieve longer time-range simultaneously keeping total displacement small compared to the size of the embryo). Importantly, we noted that the affected region (showing intense staining in the red channel) translated largely without deforming when actuated by the ferrofluid droplet, suggesting that the effective Young’s modulus of the tissue was noticeably increased by the treatment.

**FIG. S5.**
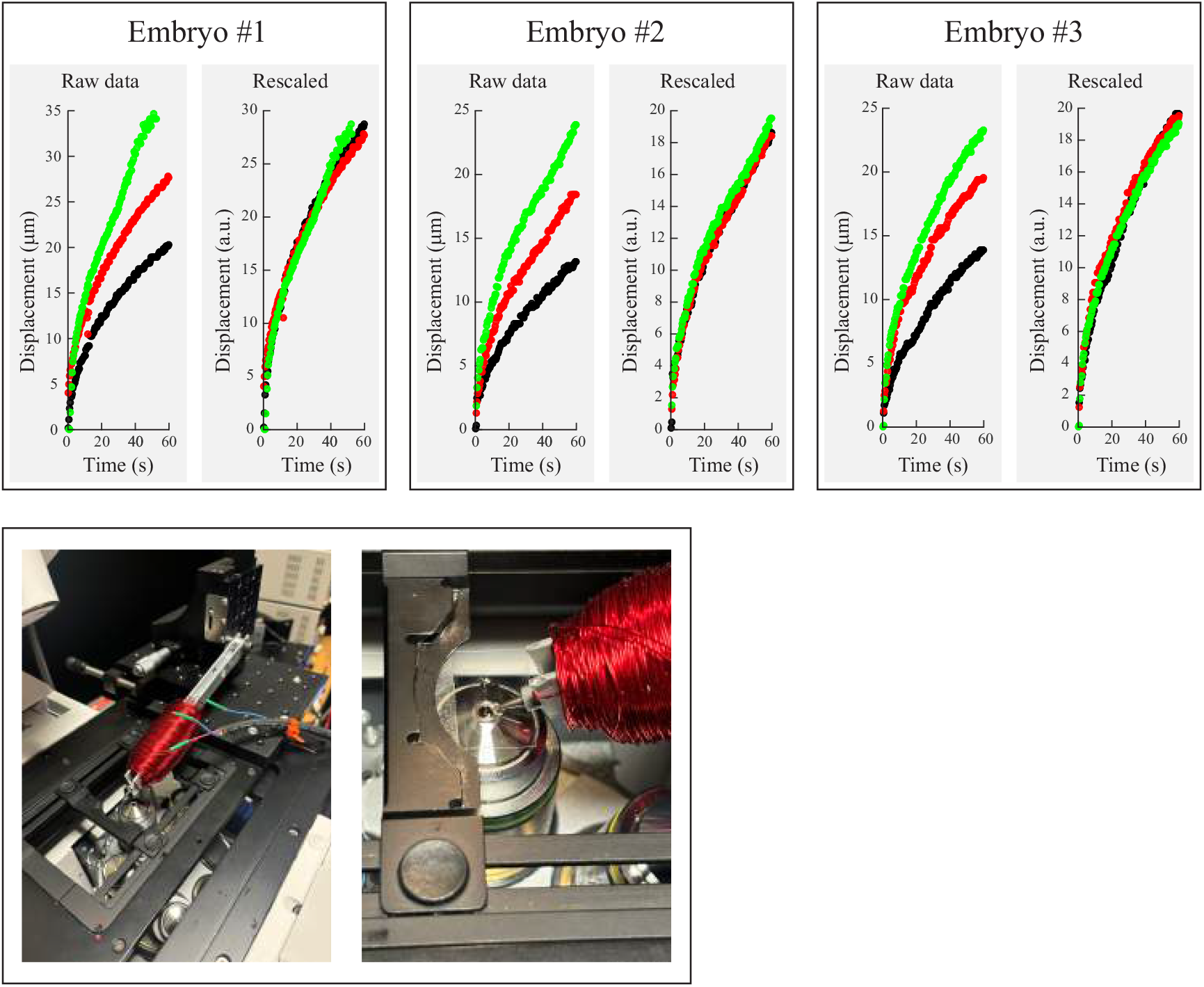
Dependence of tissue response on external force. Top panel: experimental measurements. For each embryo, three consecutive experiments were done with sufficient time for tissue relaxation in between. The current was dialed to 1, 2, or 3 A corresponding to the black, the red, and the green traces respectively. Panels on the left show traces re-scaled by the square root of the current. Bottom panel shows images of electromagnet setup. The magnet coil was made from 0.5 mm thick enameled copper wire wound around an aluminum U-bracket with outer dimensions of 12.8×10 mm and a thickness of 1.6 mm. The outer dimensions of the coil were 80 mm (length) and 33 mm (diameter). The coil comprised two roughly equal windings, that could each be powered separately, but only one of those was powered in our reported measurements. The core was made from a nail of soft iron having a length of 10 cm and a diameter of 5 mm, machined down to a narrow tip at one end. The core could be held in place within the U-bracket by a small screw in the bracket wall.

To interpret our measurements, we next turned to simulations. As may be seen from Fig. S6c, blocking turnover within a simulated circular patch of radius *R* = 85, the time-course of the point being pulled was barely affected (compare solid olive curve representing untreated case to black dotted curve representing case with blocked turnover in a large patch, in Fig. S6c). Increasing the spring constant by a factor of 10 and 100 within the affected patch changes the prefactor of the recovered power law, but not the exponent (green and blue dotted lines in Fig. S6c). Finally, blocking turnover in the entire embryo turns the power law scaling into exponential relaxation (red dotted line in Fig. S6c). In summary, while we were not able to block turnover of actin experimentally within the entire embryo for technical reasons, our experiments where turnover was blocked within a large tissue patch do agree with theoretical predictions.

**FIG. S6.**
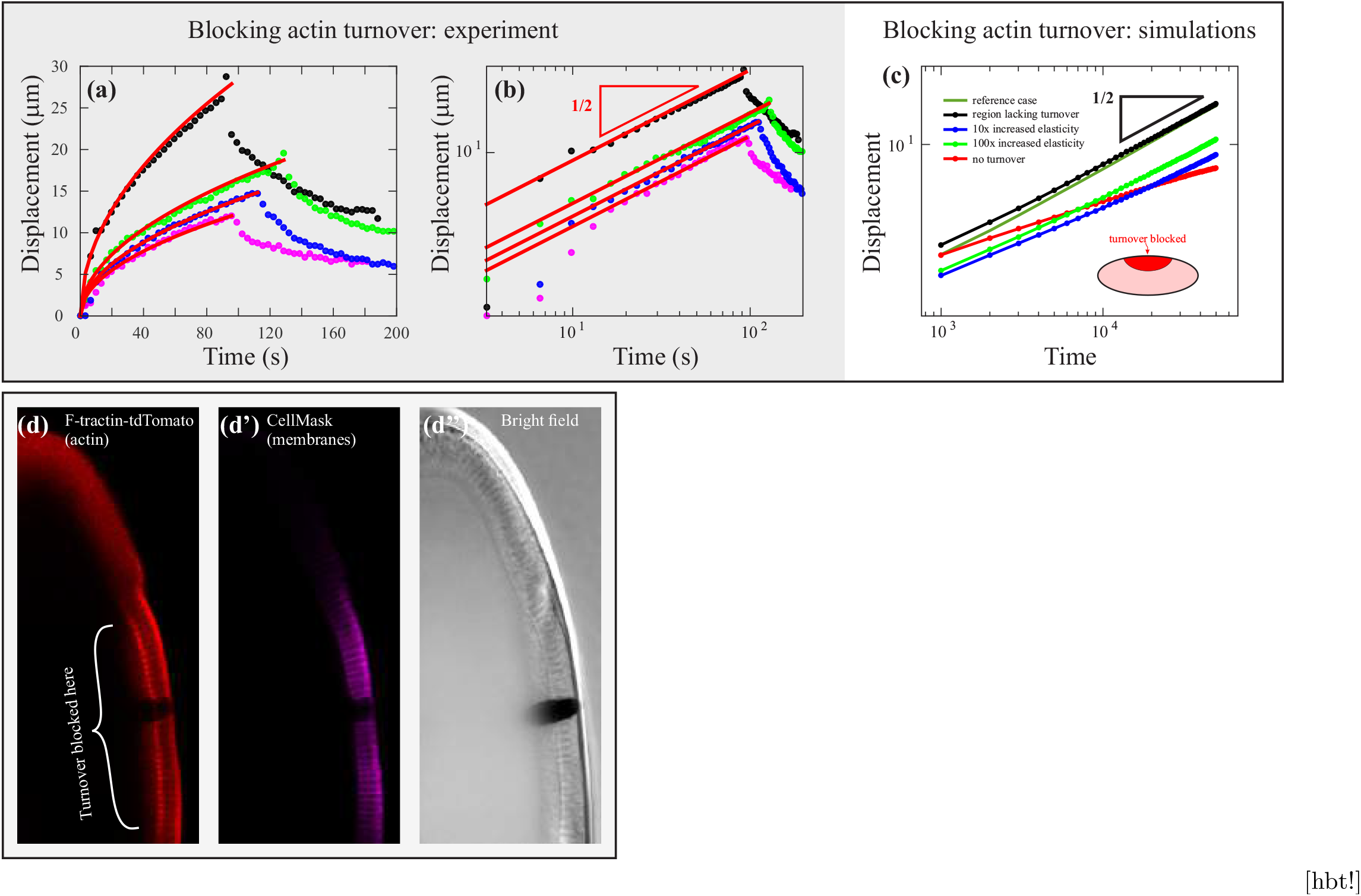
Blocking actin turnover within an extended patch of the tissue. a). Measurement data from four embryos injected with pharmacological cocktail. Cocktail was prepared by mixing 0.4 *μ*l of Deep Red CellMask (diluted 1/10 in water), and 3.6 *μ*l of jasplakinolide (800 mM solution prepared by mixing 1 *μ*l of stock solution with 2.6 *μ*l of DMSO). b). Same as middle plot, on log-log scale. c). Simulations showing the effects of blocking filament turnover within a large circular patch centered on the point where force is being applied. Olive solid curve is the original simulation with equal rate of turnover throughout the tissue. Parameters are as follows: *k*Δ*/F* = 4, *η*Δ*/*(*Fτ*) = 20, ellipsoid half-axes *R*_1_*/*Δ and *R*_2_*/*Δ = *R*_3_*/*Δ are 63 and 22. d)-d”) representative image showing a portion of an injected embryo. Red: F-tractin-tdTomato labeling polymeric F-actin. Bright red region is severely affected. Magenta: CellMask label of plasma membranes. Bright field image is shown on the right. Note that membranes are shorter in the treated region indicating that membranes are failing to extend, validating the activity of the pharmacological cocktail.

## Notes

https://github.com/mcheikh1991/Results_Scaling_Law_For_Epithelial_Tissue_Rhelogy

### Competing Interest Statement

The authors have declared no competing interest.

### Summary of Updates

Additional data included. New data concern the effects of pharmacological perturbations, the influence of finite size effects, and the scaling of the response with the external force.

## References

[1] R. Farhadifar, J. C. Roper, B. Aigouy, S. Eaton, and F. Julicher, The influence of cell mechanics, cell-cell interactions, and proliferation on epithelial packing, Curr Biol 17, 2095 (2007).

[2] S. Kim, M. Pochitaloff, G. A. Stooke-Vaughan, and O. Campas, Embryonic tissues as active foams, Nat Phys 17, 859 (2021).

[3] X. Wang, M. Merkel, L. B. Sutter, G. Erdemci-Tandogan, M. L. Manning, and K. E. Kasza, Anisotropy links cell shapes to tissue flow during convergent extension, Proc Natl Acad Sci U S A 117, 13541 (2020).

[4] M. Saadaoui, D. Rocancourt, J. Roussel, F. Corson, and J. Gros, A tensile ring drives tissue flows to shape the gastrulating amniote embryo, Science 367, 453 (2020).

[5] S. Alt, P. Ganguly, and G. Salbreux, Vertex models: from cell mechanics to tissue morphogenesis, Philos Trans R Soc Lond B Biol Sci 372, 10.1098/rstb.2015.0520 (2017).

[6] A. G. Fletcher, M. Osterfield, R. E. Baker, and S. Y. Shvartsman, Vertex models of epithelial morphogenesis, Biophys J 106, 2291 (2014).

[7] A. Hocevar and P. Ziherl, Degenerate polygonal tilings in simple animal tissues, Phys Rev E Stat Nonlin Soft Matter Phys 80, 011904 (2009).

[8] M. Rauzi, A. Hocevar Brezavscek, P. Ziherl, and M. Leptin, Physical models of mesoderm invagination in drosophila embryo, Biophys J 105, 3 (2013).

[9] T. E. Vanderleest, C. M. Smits, Y. Xie, C. E. Jewett, J. T. Blankenship, and D. Loerke, Vertex sliding drives intercalation by radial coupling of adhesion and actomyosin networks during drosophila germband extension, Elife 7, 10.7554/eLife.34586 (2018).

[10] D. P. Kiehart, J. M. Crawford, A. Aristotelous, S. Venakides, and G. S. Edwards, Cell sheet morphogenesis: dorsal closure in drosophila melanogaster as a model system, Annual review of cell and developmental biology 33, 169 (2017).

[11] Q. Wang, J. J. Feng, and L. M. Pismen, A cell-level biomechanical model of drosophila dorsal closure, Biophysical journal 103, 2265 (2012).

[12] R. Clement, B. Dehapiot, C. Collinet, T. Lecuit, and P. F. Lenne, Viscoelastic dissipation stabilizes cell shape changes during tissue morphogenesis, Curr Biol 27, 3132 (2017).

[13] K. Bambardekar, R. Clement, O. Blanc, C. Chardes, and P. F. Lenne, Direct laser manipulation reveals the mechanics of cell contacts in vivo, Proc Natl Acad Sci U S A 112, 1416 (2015).

[14] K. Nishizawa, S. Z. Lin, C. Chardes, J. F. Rupprecht, and P. F. Lenne, Two-point optical manipulation reveals mechanosensitive remodeling of cell-cell contacts in vivo, Proc Natl Acad Sci U S A 120, e2212389120 (2023).

[15] B. He, K. Doubrovinski, O. Polyakov, and E. Wieschaus, Apical constriction drives tissue-scale hydrodynamic flow to mediate cell elongation, Nature 508, 392 (2014).

[16] K. Doubrovinski, M. Swan, O. Polyakov, and E. F. Wieschaus, Measurement of cortical elasticity in drosophila melanogaster embryos using ferrofluids, Proc Natl Acad Sci U S A 114, 1051 (2017).

[17] M. I. Cheikh, J. Tchoufag, M. Osterfield, K. Dean, S. Bhaduri, C. Zhang, K. K. Mandadapu, and K. Doubrovinski, A comprehensive model of drosophila epithelium reveals the role of embryo geometry and cell topology in mechanical responses, Elife 12, 10.7554/eLife.85569 (2023).

[18] A. D’Angelo, K. Dierkes, C. Carolis, G. Salbreux, and J. Solon, In vivo force application reveals a fast tissue softening and external friction increase during early embryogenesis, Curr Biol 29, 1564 (2019).

[19] A. N. Goldner, S. M. Fessehaye, N. Rodriguez, K. A. Mapes, M. Osterfield, and K. Doubrovinski, Evidence that tissue recoil in the early drosophila embryo is a passive not active process, Mol Biol Cell 34, br16 (2023).

[20] A. N. Goldner, M. I. Cheikh, M. Osterfield, and K. Doubrovinski, Viscous shear is a key force in drosophila ventral furrow morphogenesis, https://www.biorxiv.org/ 10.1101/2021.04.21.440835 (2023).

[21] N. Desprat, A. Richert, J. Simeon, and A. Asnacios, Creep function of a single living cell, Biophys J 88, 2224 (2005).

[22] B. D. Hoffman, G. Massiera, K. M. Van Citters, and J. C. Crocker, The consensus mechanics of cultured mammalian cells, Proc Natl Acad Sci U S A 103, 10259 (2006).

[23] A. D’Angelo, K. Dierkes, C. Carolis, G. Salbreux, and J. Solon, In vivo force application reveals a fast tissue softening and external friction increase during early embryogenesis, Curr Biol 29, 1564 (2019).

[24] H. G. Dobereiner, B. Dubin-Thaler, G. Giannone, H. S. Xenias, and M. P. Sheetz, Dynamic phase transitions in cell spreading, Phys Rev Lett 93, 108105 (2004).

[25] L. Figard, L. Zheng, N. Biel, Z. Xue, H. Seede, S. Coleman, I. Golding, and A. M. Sokac, Cofilin-mediated actin stress response is maladaptive in heat-stressed embryos, Cell Rep 26, 3493 (2019).

[26] H. S. Seung and D. R. Nelson, Defects in flexible membranes with crystalline order, Phys Rev A Gen Phys 38, 1005 (1988).

[27] H. Liang and L. Mahadevan, Growth, geometry, and mechanics of a blooming lily, Proc Natl Acad Sci U S A 108, 5516 (2011).

[28] M. Wyart, H. Liang, A. Kabla, and L. Mahadevan, Elasticity of floppy and stiff random networks, Phys Rev Lett 101, 215501 (2008).

[29] C. P. Broedersz, X. Mao, T. C. Lubensky, and F. C. MacKintosh, Criticality and isostaticity in fibre networks, Nature Physics 7, 983 (2011).

[30] J. T. Blankenship, S. T. Backovic, J. S. Sanny, O. Weitz, and J. A. Zallen, Multicellular rosette formation links planar cell polarity to tissue morphogenesis, Dev Cell 11, 459 (2006).

## References

[1] M. I. Cheikh, J. Tchoufag, M. Osterfield, K. Dean, S. Bhaduri, C. Zhang, K. K. Mandadapu, and K. Doubrovinski, A comprehensive model of drosophila epithelium reveals the role of embryo geometry and cell topology in mechanical responses, Elife 12, 10.7554/eLife.85569 (2023).

[2] K. Doubrovinski, M. Swan, O. Polyakov, and E. F. Wieschaus, Measurement of cortical elasticity in drosophila melanogaster embryos using ferrofluids, Proc Natl Acad Sci U S A 114, 1051 (2017).

[3] G. E. Peng, S. R. Wilson, and O. D. Weiner, A pharmacological cocktail for arresting actin dynamics in living cells, Molecular biology of the cell 22, 3986 (2011).

[4] C. A. Wilson, M. A. Tsuchida, G. M. Allen, E. L. Barnhart, K. T. Applegate, P. T. Yam, L. Ji, K. Keren, G. Danuser, and J. A. Theriot, Myosin ii contributes to cell-scale actin network treadmilling through network disassembly, Nature 465, 373 (2010).

[5] A. J. Spracklen, T. N. Fagan, K. E. Lovander, and T. L. Tootle, The pros and cons of common actin labeling tools for visualizing actin dynamics during drosophila oogenesis, Developmental biology 393, 209 (2014).

